# Profiling Antibiotic Resistance Determinants in Ancient Permafrost Microbiomes

**DOI:** 10.1101/2024.05.20.595077

**Authors:** Sankaranarayanan Gomathinayagam, Gothandam Kodiveri Muthukaliannan

## Abstract

Antimicrobial resistance presents a formidable challenge, yet its existence predates the introduction of antibiotics. Our study delves into the presence of antimicrobial resistance determinants (ARDs) in ancient permafrost microbiomes, comparing them with contemporary soil and pristine environments. Majority of the samples are from regions around Beringia, encompassing parts of Russia and Alaska, with only one sample originating from the Tien Shan Mountain range in Kyrgyzstan. From over 2.3 tera base pairs of raw metagenomic data, we assembled about 1.3 billion metagenomic contigs and explored the prevalence of ARDs in them. Our findings reveal a diverse array of ARDs in ancient microbiomes, akin to contemporary counterparts. On average, we identified 2 ARDs per ribosomal protein gene in ancient samples. *Actinomycetota, Bacillota*, and several thermophiles were prominent carriers of ARDs in Chukochi and Kamchatkan samples. Conversely, ancient permafrost from the Tien Shan Mountain range exhibited no Thermophiles or *Actinomycetota* carrying ARDs. Both ancient and contemporary microbiomes showcased numerous divergent ARDs, with approximately 40% identity to genes in antibiotic resistance gene databases. Antibiotic inactivation-type ARDs exhibited purifying selection with contemporary resistance genes, as estimated by dN/dS ratio. Importantly, we retrieved 359 putative complete viruses from ancient microbiomes and none of them harboured any ARDs.

## Introduction

Microorganisms, as some of the oldest extant organisms, have endured myriad challenging environments over eons, predating significant events such as the Great Oxygenation Event and the subsequent oxygenated world. They have demonstrated remarkable resilience and adaptability, thriving for billions of years. Throughout their extensive history, microorganisms have encountered and adapted to numerous toxic environments, necessitating the development of mechanisms to mitigate such challenges. For instance, the transition to an oxygenated environment posed a threat to anaerobic microorganisms, compelling them to evolve antioxidant mechanisms and other defensive strategies against environmental toxins.

In a similar vein, the phenomenon of antimicrobial resistance transcends time, likely originating alongside the production of antimicrobials by microorganisms themselves. Even preceding the discovery and widespread use of antibiotics, evidence suggests the presence of antimicrobial resistance within microbial populations[1]. This antimicrobial synthesis and development of resistance is thought to have arisen as a result of competition within their ecological niches, illustrating the ancient and ongoing arms race between microorganisms and antimicrobial substances. But it would be interesting to investigate what the scenario of the antimicrobial resistome must have been in the microbial environment before the anthropocene, in contrast to the age of extensive antibiotic use.

There are few historical collections of bacterial cultures dating back to the time antibiotics were discovered, such as the collection assembled by E.G.D. Murray spanning the period from 1917 to 1954. This collection is known to still be maintained at the National Collection of Type Cultures, Public Health England. Collections like these would provide valuable insights into microbes from the time of clinical introduction of antibiotics[2].

Furthermore, studies examining historical microbiomes, such as those derived from dental calculus of ancient humans and animals, coprolites, mummified bones etc., offer another window into the past. In a previous study, we investigated antimicrobial resistance genes in Neanderthal dental calculus and coprolite microbiomes. However, we encountered challenges related to post-mortem DNA damage, which impeded the retrieval of complete antimicrobial resistance determinants (ARDs) from these ancient microbiomes. Additionally, contamination from soil and other environmental sources posed further obstacles to the accurate detection of ancient antimicrobial resistance genes distinct from their modern counterparts. Moreover, the authenticity of the identified putative ARDs could not be confirmed through cytosine deamination (typical characteristic of ancient DNA fragment) at the ends of the reads corresponding to the respective ARDs.

Similarly, some researchers have explored ancient permafrost from Arctic and Antarctic regions, uncovering antimicrobial resistance genes from pre-historic era. However, some of these approaches utilized targeted amplicon methods with primers designed for modern-day antimicrobial resistance genes, potentially leading to underestimation or overestimation of the true quantitative presence of antimicrobial resistance genes. Moreover, such approaches may potentially introduce PCR bias, obscuring the authentic nucleotide sequences of ancient antimicrobial resistance genes[1]. To address these challenges, we aimed to comprehensively explore ancient permafrost microbiomes and profile the entire resistome without employing any biased or targeted approaches. It is important to note that ancient antimicrobial resistance genes may not necessarily have representatives in present-day antibiotic resistance gene databases. These pristine environments offer undisturbed settings for studying microbial environments in the context of antimicrobial resistance with reduced post mortem DNA damage.

Earlier studies have suggested that antimicrobial resistance genes retrieved from ancient microbiomes are “divergent” from contemporary counterparts[3]. Building on these findings, we sought to address several key questions regarding ancient antimicrobial resistance genes. These include how divergent are antimicrobial resistance genes from palaeomicrobiomes from their modern counterparts, whether modern-day antimicrobial resistance genes, particularly after the introduction of antibiotics, have undergone positive evolutionary changes to become more effective against antibiotics, and whether there are differences in their collective modes of action. Additionally, we aimed to investigate whether bacteriophages generally carry antimicrobial resistance genes in palaeomicrobiological niches or if only a few of them harbour such genes due to selection pressure resulting from the irrational use of antibiotics in modern times.

Our study entails a comprehensive analysis of ancient microbiomes from existing genomic databases, profiling ARDs, identifying conserved domains relevant for antimicrobial resistance, and analysing selection pressure by estimating the Ka/Ks value. Furthermore, we aimed to meticulously classify potential complete bacteriophages and identify ARDs(if at all present) within them.

## Methods

### Data collection

Ancient metagenome datasets were systematically searched in the NCBI BioProject and community-curated SPAAM AncientMetagenomeDir using specific keywords to identify relevant studies. The keywords employed for the search included “Permafrost AND ancient Metagenome,” “Icecore and Metagenome,” and “Cryosoil AND metagenome.” Additionally, a taxonomy search with the taxon ID for permafrost metagenome ‘1082480’ was conducted.

The initial search yielded a total of 1427 Sequence Read Archives across 116 BioProjects upto June 2023. Subsequently, these projects underwent further scrutiny, specifically excluding targeted loci methods such as the 16S amplicon method. The projects in which the antiquity of the sample was not clearly established were also ignored. As a result, the final number of projects was narrowed down to a total of ten, namely PRJEB47746 [4], PRJNA266334 [5], PRJNA343018 [6], PRJNA350710 [7], PRJNA505516 [8], PRJNA596250 [9], PRJNA601698 [10], PRJNA680161 [11], PRJNA438924 [12], and PRJDB5557 [13]. The age of the samples in these projects ranged from 7000 years ago to 1100 kilo years ago (Supplementary Table S1).

### Geographical distribution of source data

The above-mentioned studies were conducted in geographical locations primarily concentrated in Russia, with only three of them located outside Russia as shown in Figure 1. These include two projects from Alaska, one focusing on Fox, renowned for cold regions research, and the other on the Grigoriev Glacier in the Tien Shan Mountain range of Kyrgyzstan, Central Asia. Other samples were obtained from diverse geographical locations within Russia, such as Kamchatka, a volcanic peninsula in the Russian Far East; the Kolyma-Indigirka Lowland, characterized by extensive permafrost in northeastern Siberia, Russia; regions such as Omolon, Gydan, and Bykovsky in Siberia, distinguished by their geological formations; specific sites along the Alazeya River within the Kolyma-Indigirka Lowland; geological formations like the Yedoma and Olyor suites within the same region; and areas encompassing the Alazeya River and Cape Chukochya. Additionally, Kon’kovaya which represent specific landmarks or formations within the Kolyma-Indigirka Lowland. The sampling methodologies employed in these studies follow a consistent approach aimed at minimising contamination during sample collection and processing. Dry drilling techniques were universally utilised, with no drilling fluid employed to avoid introducing contaminants into the permafrost samples. Core segments were extracted using air pressure or power drills, avoiding contamination from the surrounding environment. Additionally, in some cases, the outer peripheral layer of the cores was removed using a sterile knife to eliminate potential surface contaminants. The inner part of the cores, which is less likely to be affected by surface contamination, was targeted for sampling. Furthermore, in one study, the outer layer was coated with a control organism and then scraped off with a sterile knife to ensure thorough removal of any potential contaminants. DNA isolation has been carried out in clean room setups, with stringent measures to prevent cross-contamination.

**Figure 1.**
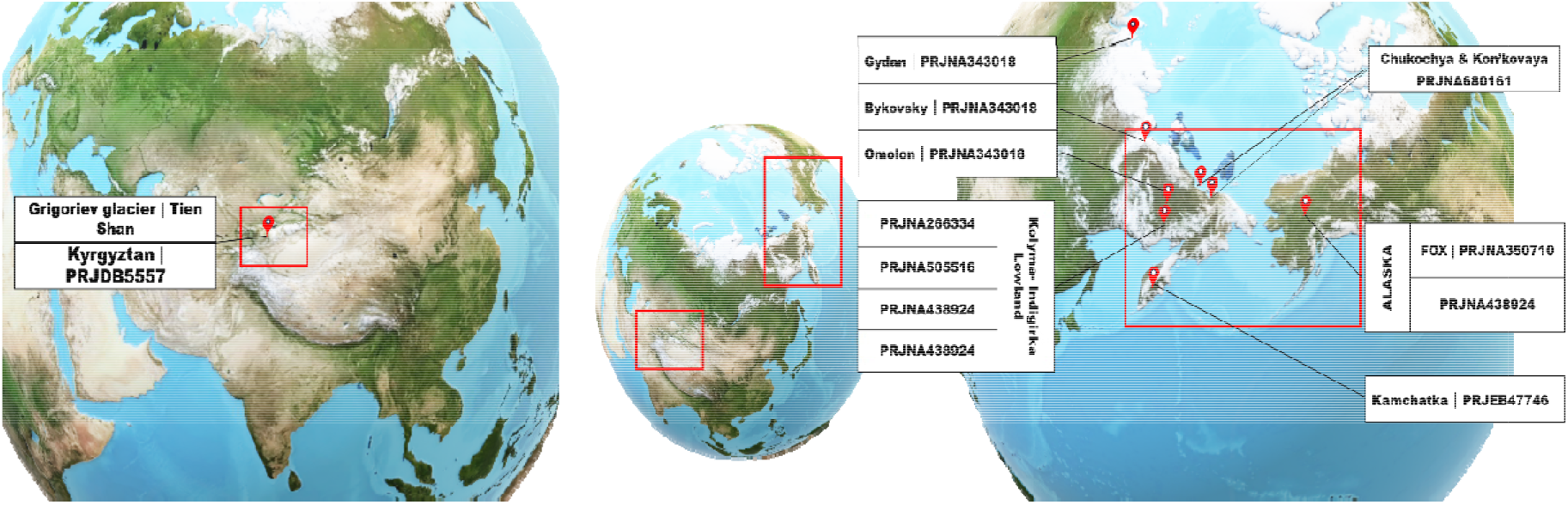
Geographical locations of studied samples across different projects.

Some projects, like PRJNA505516 and PRJNA680161, aimed to identify thriving microbes in ice cores using LIVE/DEAD staining. They distinguished the isolated genome from thriving microbes (based on LIVE/DEAD staining) as the intracellular genome, while the DNA from debris microbes was categorized as extracellular or environmental DNA (Supplementary Data S1). In PRJNA680161, replicates of samples have been prepared, with one set undergoing DNA damage repair using the preCR DNA repair cocktail, while the other set retained native DNA. Only the replicate with repaired DNA damage was selected for this study. In the project PRJNA438924, samples from both the Pleistocene and Holocene epochs were examined. However, only samples dating back to the Pleistocene were selected for inclusion in this study.

### Control metagenome data

As control samples, we selected three additional studies, apart from three contemporary samples analysed in study PRJEB47746. One of these studies examined metagenomes sequenced from anoxic sites, including permafrost characterized by low water availability, low water activity, low temperature, restricted availability of nutrients, and acidic conditions - PRJEB28336. Other two are metagenome samples from municipal waste dumps/landfills - PRJNA718480 and PRJNA606662.

### Profiling ancient ARDs in metagenomes

The paired-end FASTQ files were imported into the KBase^®^ server using the Import Paired-End Files utility [14]. Subsequently, the quality of the FASTQ files was assessed for the presence of adapters and read fragmentation using FASTQC tool. Adapters were removed using the Trimmomatic[15] tool, resulting in clean FASTQ files. These clean files were employed for metagenome assembly using the MEGAHIT tool in ‘meta large’ mode, with a minimum contig length threshold set at 1000 bps [16].

The assembled contigs were downloaded onto a Linux workstation, and the AMRFinderPlus tool was executed with a minimum coverage of 0.90 and minimum identity threshold of 0.40 [17]. Hits containing internal stop codons and those originating from partial gene fragments (PARTIALX) were filtered out. The nucleotide sequences of the remaining hits were then translated using the Transeq [18] command line tool. Next, the translated protein sequences underwent a search for conserved domains using InterPro [19].

For ARD count normalisation, the number of identifiable 5S, 16S, and 23S genes from the assembled contigs was analysed using the Barrnap tool with default settings [20]. The absolute count of these genes was utilised to normalise the absolute count of ARDs obtained from the AMRFinderPlus run.

### Selection pressure estimation

The Ka/Ks ratio was computed for ORFs paired with their respective reference genes using Ka_Ks Calculator Tool version 3 [21]. Protein sequences underwent alignment using the MAFFT aligner [22]. This alignment data was utilized for codon alignment of nucleotide sequences employing PAL2NAL with the “-nogap” argument, aimed at removing gap lengths not divisible by 3 and non-overlapping regions [23]. Subsequently, the resulting clustal-formatted, codon-informed nucleotide alignment was transformed into an AXT file. This AXT file was then provided as input to the Ka_Ks Calculator, and the analysis was conducted using model-averaged method to estimate the selection pressure between the sequences.

### Mapping based assembly

For samples that didn’t yield any assembled contigs, a k-mer-based read-level ARD finding approach was implemented using KMA tools [24]. Initially, the CARD-ARG database and NCBI’s Refgene catalogue for Antibiotic Resistance Genes (v2020.10.12) were merged [25], and a KMA index was generated. Subsequently, the KMA aligner was employed to align the reads to this index, and the output was rendered as a SAM format file. Similarly, the GTDB SSU (bac120) database was indexed, and the reads were aligned to this index to identify SSU (Small Subunit) genes for ARD count normalization [26]. A mapping coverage threshold of 0.70 was selected for both ARDs and SSUs from the SAM files.

### Taxonomic classification of ARD contigs

For rapid taxonomical identification of ARD bearing contigs, Kraken2 tool was employed to analyse the contigs against the Kraken2 Standard-8 microbial database (https://genome-idx.s3.amazonaws.com/kraken/k2_standard_08gb_20240112.tar.gz accessed on Feb 12, 2024) with default parameters [27]. The alpha diversity indices were then estimated using the Paleontological Software Tool (PAST v4) [28].

### Classification and retrieval of complete viral genomes

To classify viral contigs from the metagenome-assembled contigs, a hierarchical classification method was employed. Initially, the contigs were superficially classified using a deep learning method by DeepVirFinder, with a contig length cutoff of 3000 bp. Contigs with a viral score of at least 0.7 were selected for further classification [29]. Subsequently, the putative viral contigs underwent classification by VIBRANT version 1.2.1, which identifies viral hallmark proteins in the contigs [30]. Contigs classified as complete-circular by VIBRANT were then retrieved and subjected to AMRFinderPlus analysis to profile ARDs in the complete viruses with a minimum identity of 0.40 and a minimum coverage of 0.40.

## Results

### Profiling of ARDs in ancient microbiome

A total of approximately 2.3 tera basepairs of data were assembled into contigs using MEGAHIT, resulting in 13,919,982 contigs (>1000 bps) in total. Three samples from the project PRJNA505516 and one sample from project PRJNA596250 did not yield any contigs (see Table 3). Meanwhile, the samples from project PRJNA266334 yielded the lowest number of contigs. Among all the ancient metagenome contigs, AMRFinderPlus identified a total of 33,481 ARDs, while Barrnap version 0.7 identified 16,541 ribosomal RNA genes (5S, 16S, and 23S). This translates to roughly 2 ARDs per ribosomal gene identified in the contigs. Notably, in one sample (PRJDB5557: DRR088405), 6 ARDs were identified, but no ribosomal genes could be detected. Table 1 illustrates that ARDs ranged as high as 3.25±0.565 per ribosomal housekeeping gene in the project PRJNA47746, with the lowest occurrence observed in project PRJNA596250 (0.20±0.028). In contrast, in contemporary samples from project PRJNA47746, the occurrence was 0.89; in PRJNA718480, it was 1.91; and in PRJNA606662, it was 0.44. The identified classes of resistance genes encompassed beta-lactam, tetracycline, macrolides including phenicols, fluoroquinolones, among others.

**Table 1:** Survey of ARDs present in metagenome of various ancient samples.

| **Project accession** | **Total contigs count<sup>a</sup>** | **Total ARD count<sup>b</sup>** | **Total 5s, 16s, 23s count<sup>c</sup>** | **Mean RSU normalised ARD count ± SEM** |
| --- | --- | --- | --- | --- |
| PRJNA47746 | 83,36,212 | 24,851 | 9,613 | 3.25±0.565 |
| PRJNA266334 | 76 | 0 | 0 | 0 |
| PRJNA680161 | 4,78,607 | 768 | 954 | 0.82±0.065 |
| PRJNA601698 | 6,43,806 | 1,166 | 1,356 | 1.23±0.405 |
| PRJNA350710 | 5,01,690 | 887 | 1,135 | 0.59±0.153 |
| PRJNA438924 | 11,99,868 | 4,140 | 1708 | 2.66±0.359 |
| PRJNA596250 | 90,014 | 56 | 273 | 0.20±0.028 |
| PRJNA505516 | 4,40,412 | 450 | 697 | 0.43±0.113 |
| PRJDB5557 | 42,788 | 141 | 60 | 1.13±0.479 |
| PRJNA343018 | 13,39,308 | 1,022 | 745 | 1.24±0.228 |

a – Total number of contigs above 1000 bps length across different samples; b – Absolute number of ARDs in the assembled contigs with > 90% of coverage and > 0.40% of identity to the reference sequences in AMRFinder database; c – detected number of 5s, 16s, 23s ribosomal subunit genes detected by barnap tool.

**Table 2:**
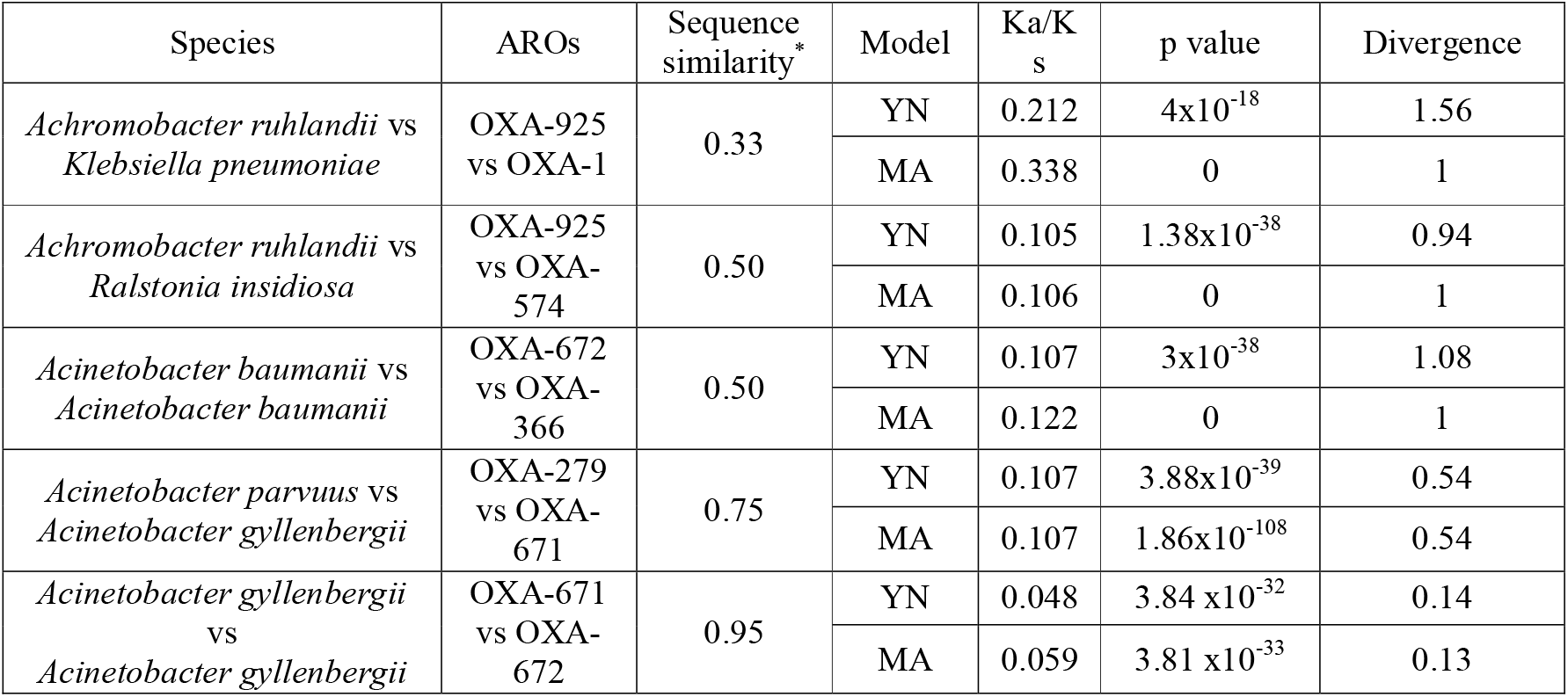
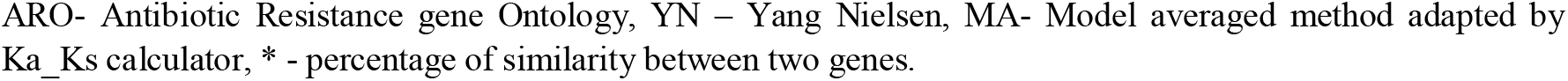
Validation of model selection for Ka/Ks estimation between ancient and contemporary ARDs.

| Species | AROs | Sequence similarity* | Model | Ka/Ks | p value | Divergence |
| --- | --- | --- | --- | --- | --- | --- |
| *Achromobacter ruhlandii* vs *Klebsiella pneumoniae* | OXA-925 vs OXA-1 | 0.33 | YN | 0.212 | $4 \times 10^{-18}$ | 1.56 |
|  |  |  | MA | 0.338 | 0 | 1 |
| *Achromobacter ruhlandii* vs *Ralstonia insidiosa* | OXA-925 vs OXA-574 | 0.50 | YN | 0.105 | $1.38 \times 10^{-38}$ | 0.94 |
|  |  |  | MA | 0.106 | 0 | 1 |
| *Acinetobacter baumannii* vs *Acinetobacter baumannii* | OXA-672 vs OXA-366 | 0.50 | YN | 0.107 | $3 \times 10^{-38}$ | 1.08 |
|  |  |  | MA | 0.122 | 0 | 1 |
| *Acinetobacter parvuus* vs *Acinetobacter gyllenbergii* | OXA-279 vs OXA-671 | 0.75 | YN | 0.107 | $3.88 \times 10^{-39}$ | 0.54 |
| | | | MA | 0.107 | $1.86 \times 10^{-108}$ | 0.54 |
| *Acinetobacter gyllenbergii* vs *Acinetobacter gyllenbergii* | OXA-671 vs OXA-672 | 0.95 | YN | 0.048 | $3.84 \times 10^{-32}$ | 0.14 |
| | | | MA | 0.059 | $3.81 \times 10^{-33}$ | 0.13 |

ARO- Antibiotic Resistance gene Ontology, YN – Yang Nielsen, MA- Model averaged method adapted by Ka\_Ks calculator, \* - percentage of similarity between two genes.

**Table 3:** ARDs and ribosomal proteins identified directly from the raw reads by KMA.

| Project | Accession no. | Base pairs | ARD above 0.70 | SSU above 0.70 |
| --- | --- | --- | --- | --- |
| PRJNA266334 | SRR1653578 | 91 Mbps | 0 | 5 |
|  | SRR1653579 | 91 Mbps | 0 | 1 |
| PRJNA505516 | SRR8188252<sup>i</sup> | 7.1 Gbps | 0 | 8 |
|  | SRR8188255<sup>i</sup> | 6 Gbps | 0 | 1 |
|  | SRR8188256<sup>e</sup> | 2.8 Gbps | 0 | 0 |
| PRJNA596250 | SRR13615822 | 5 Kbps | 0 | 0 |

i- DNA extracted intracellularly from probably living bacteria. e- environmental DNA from the ancient permafrost;

### Distribution of identity percentages between ARDs from ancient metagenome and contemporary metagenome

The distribution of identity percentages was analysed, considering a minimum identity threshold of 40% and a relatively stringent minimum coverage of 90%. To verify the putative identification of genes as ARGs, a conserved domain search was conducted using Interproscan. Across the samples and ages, the majority of the identified genes exhibited identity percentages between 40% and 60%, as depicted in Figure 2. Notably, samples from project PRJEB47746 displayed a relatively well-distributed range of identity percentages for ARDs compared to their counterparts. Conversely, projects PRJNA505516 and PRJNA596250 exhibited only a few ARDs with identity percentages above 60%, coinciding with a relatively lower number of identified ARDs. In contemporary metagenome projects such as PRJNA606662 and PRJNA718480, the identity of ARDs was marginally higher than in the ancient samples. ARDs found in these samples demonstrated comfortable similarity percentages of up to 70% and 90%, respectively. In contrast, ARDs from contemporary pristine samples in project PRJEB28336 clustered within 60% similarity percentages. This suggests that metagenomic ARDs, regardless of age, exhibit very low similarity to present reference ARG database.

**Figure 2.**
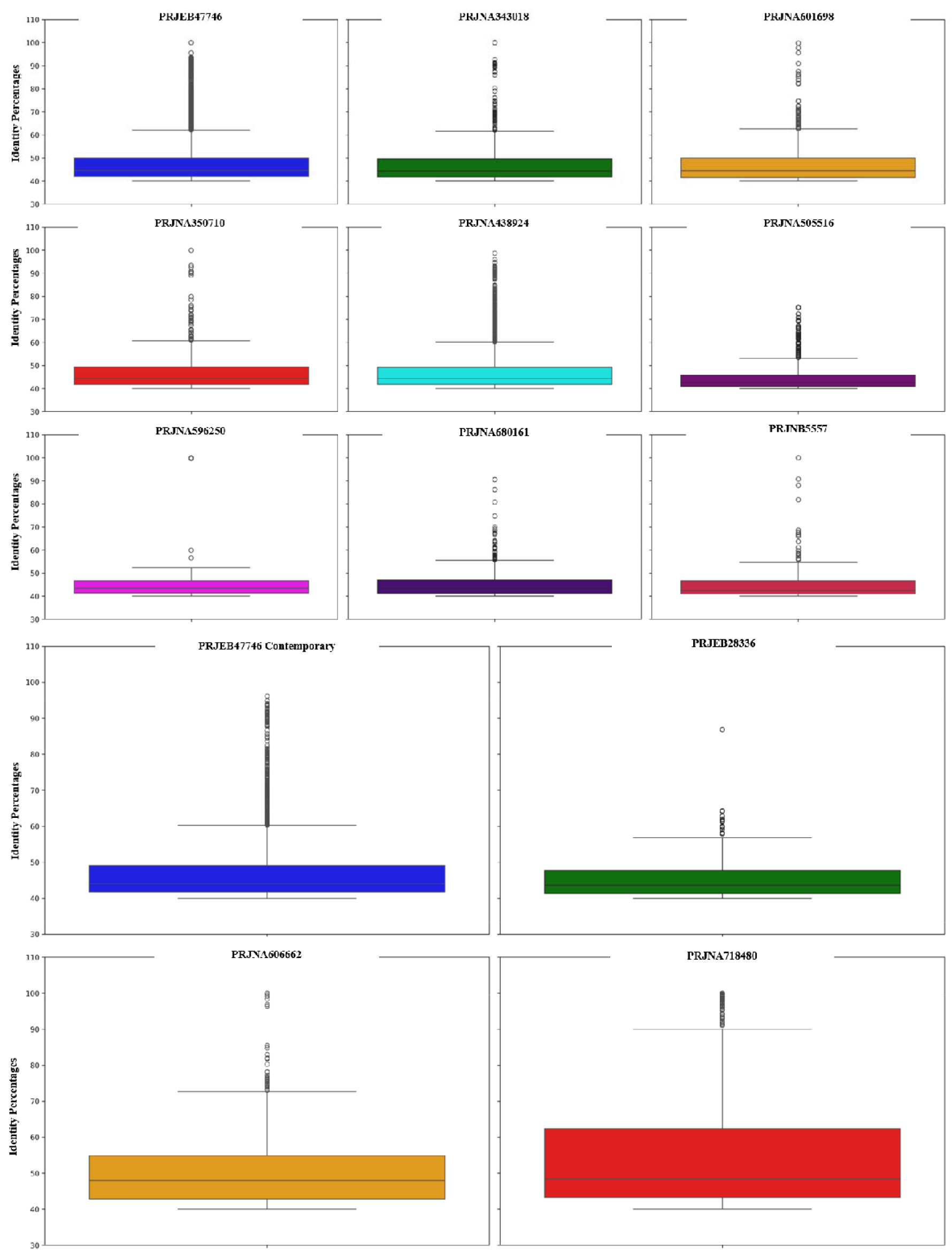
Box plot illustrating distribution of identity percentages between contemporary (Panel B) and ancient ARDs (Panel A)

### Abundance of ARDs based on resistance mechanisms and classes

The ARDs were classified into three classes based on their resistance mechanisms, such as antibiotic inactivation, antibiotic efflux, and antibiotic target protection/alteration/replacement as shown in Figure 4. In the high-depth metagenomes from the project PRJEB47746, ARDs were almost equally distributed among these three resistance mechanisms. However, in projects PRJNA596250 and PRJNA680161, ARDs related to antibiotic target protection/alteration/replacement were notably dominant. Regarding contemporary samples, pristine metagenomes exhibited a higher proportion of ARDs associated with antibiotic target protection/alteration/replacement, while antibiotic inactivation ARDs were less prominent. Conversely, samples from landfills or municipal wastes showed insignificant differences between these three classes of resistance mechanisms.

In a similar manner, the profiled ARDs were categorized into classes of antibiotics to which they confer resistance, as illustrated in Figure 3b. It can be observed that the ancient microbiome harbours resistance genes for Trimethoprim, quinolones, fosfomycin, mupirocin, nitroimidazole, sulfanamide, and other classes of antibiotics. Worth noting is that the mentioned classes of antibiotics are primarily synthetic or semi-synthetic. However, the genes conferring resistance are already ubiquitously present enzymes in bacteria, or resistance arises due to point mutations to the targets of antibiotics. For instance, resistance to Trimethoprim can occur through the inactivation of dihydrofolate reductase, while quinolone resistance may result from mutations in DNA gyrase and type IV topoisomerases. Dihydrofolate reductase, DNA gyrase and topoisomerase IV are all core essential enzymes in bacteria.

**Figure 3a:**
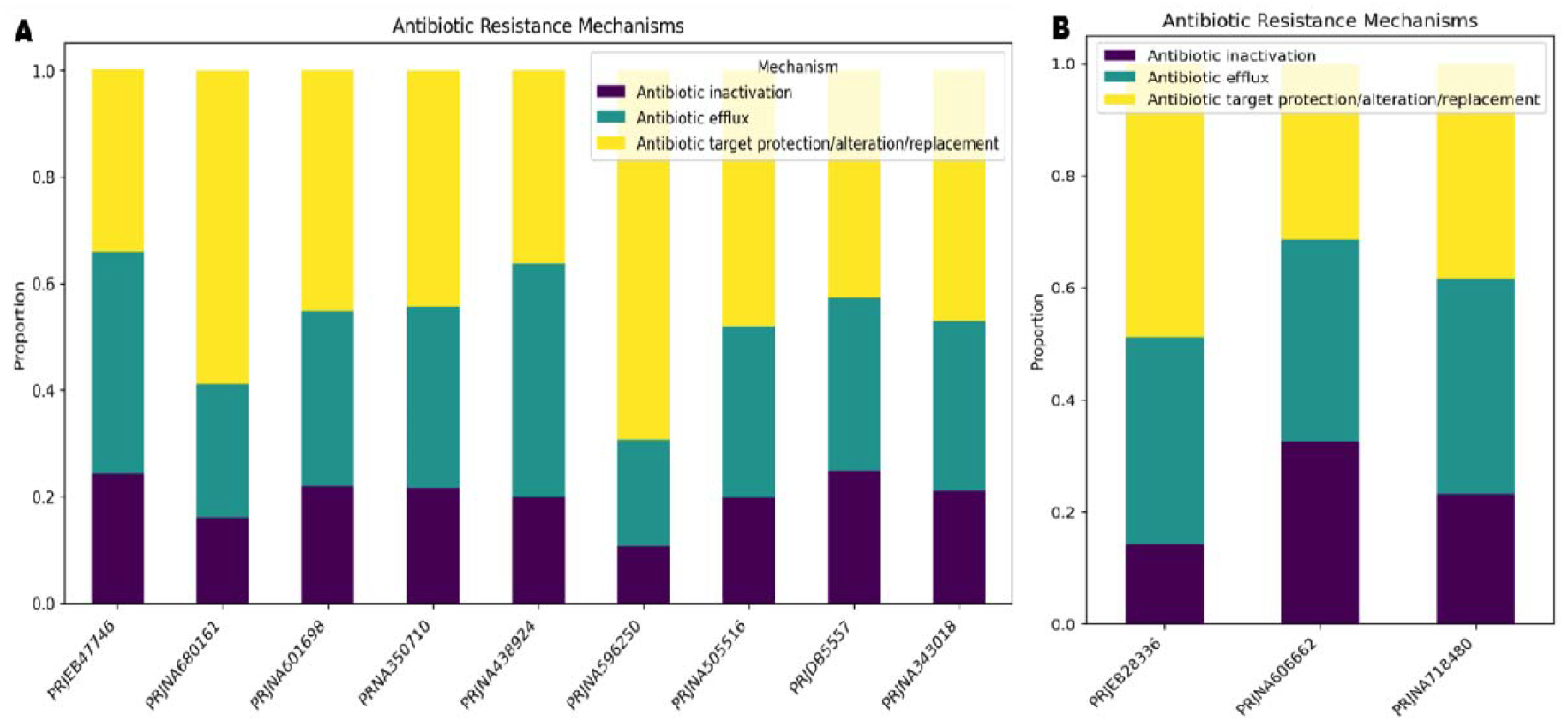
Bar plot illustrating the composition of ARDs belonging to Antibiotic inactivation, Antibiotic efflux and Antibiotic target protection/alteration/replacement in both ancient metagenomes (Tile A) and contemporary metagenomes (Tile b).

**Figure 3b:**
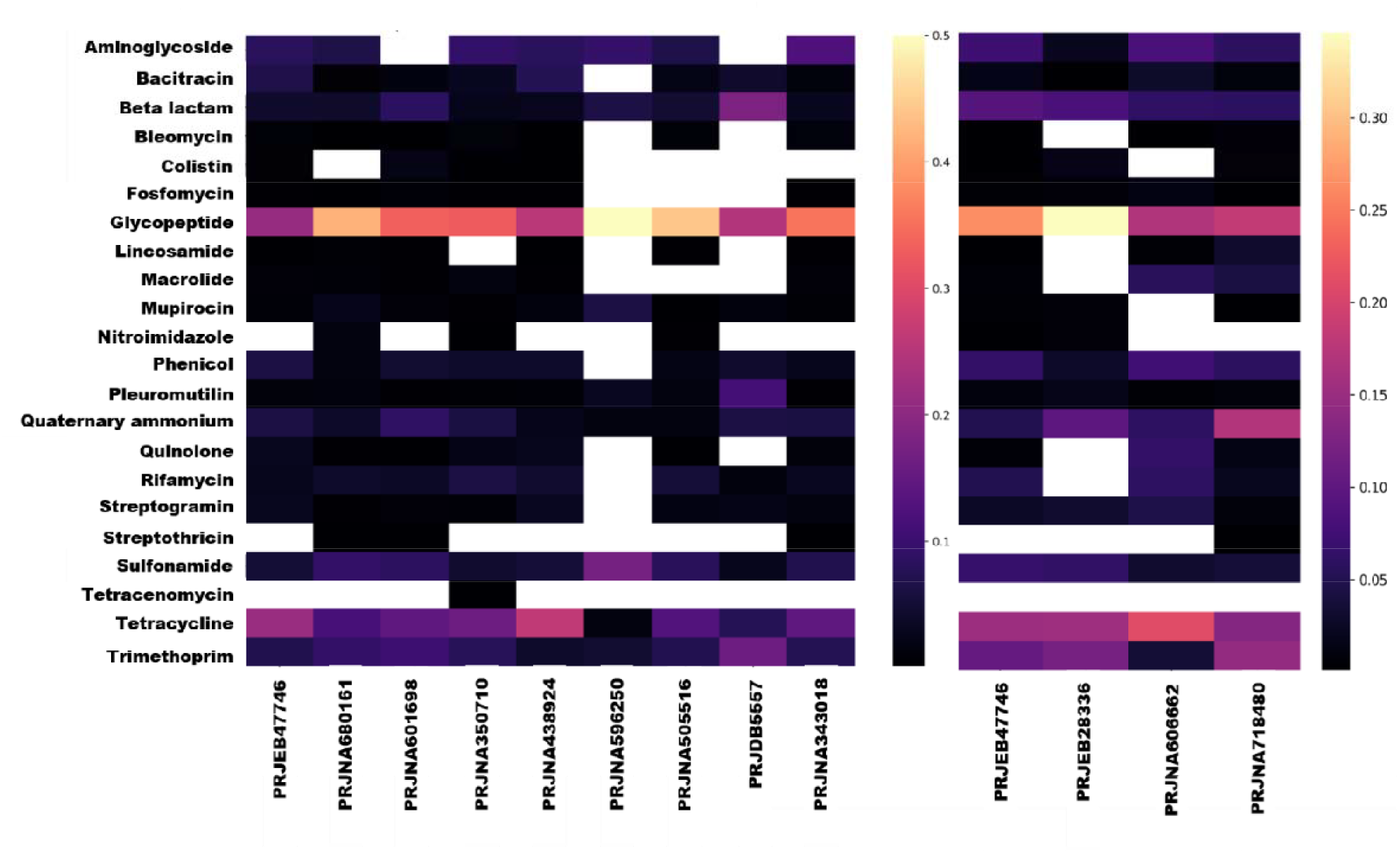
Profiled ARDs classified based on resistance to different antibiotic classes.

### Taxa of contigs carrying ARDs

A heat map was generated based on the taxonomy assignments by the Kraken tool for contigs carrying ARDs, illustrating abundance versus taxa up to the phylum level (see Figure 4). Among the studied projects, PRJEB47746 displayed a diverse range of taxa, with the majority belonging to *Pseudomonadota* and *Actinomycetota*. Interestingly, *Actinomycetota* did not exhibit the presence of ARDs in project PRJDB5557. Additionally, thermophilic phyla such as *Thermodesulfobacteriota, Deinococcota, Aquificota, Thermomicrobiota, Rhodothermota*, and *Thermoproteota* were not identified in this particular project. Following *Pseudomonadota* and *Actinomycetota, Bacillota* emerged as a consistent carrier of ARDs across all projects. Additionally, *Cyanobacteriota, Myxococcota, Bacteroidota, Gemmatimonadota, Acidobacteriota, Thermodesulfobacteriota*, and *Deinococcota* were identified as widespread carriers of ARDs across multiple projects.

**Figure 4.**
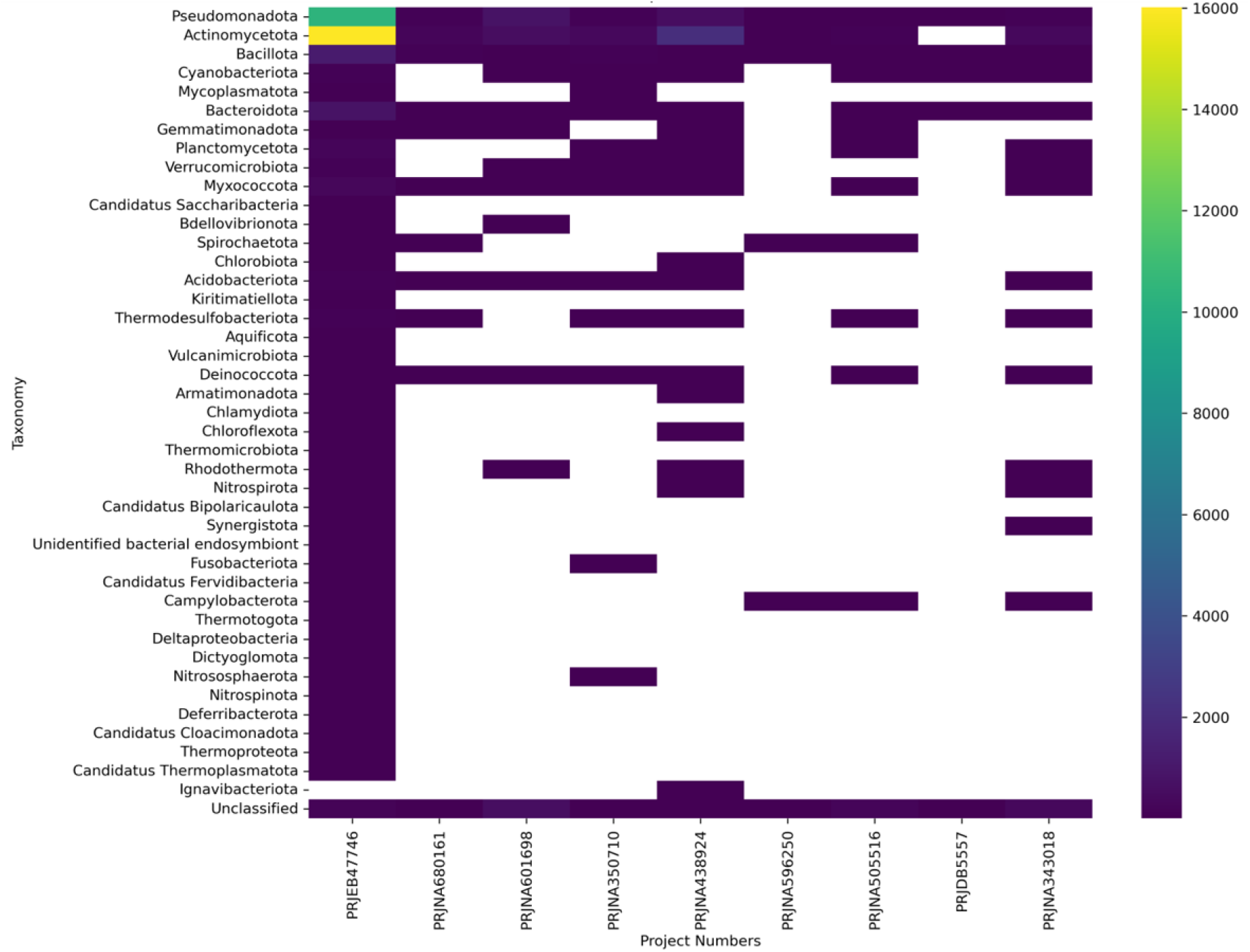
Heat map showing abundance of ARGs carrying phyla across different projects. Factors affecting ARD abundance.

We conducted a correlation analysis of the normalized ARD count with the age of samples, encompassing both contemporary and ancient samples (Supplementary Table S2). Using an inverse-ranked Spearman correlation analysis, we examined whether a decrease in age correlates with an increase in ARD count in the assembled contigs. The analysis revealed a mild negative correlation of −0.15, which was deemed statistically insignificant as it fell below the critical *t* value. Additionally, we explored whether diversity serves as a contributing factor to ARD abundance. Pearson’s correlation coefficient was employed to assess the correlation between the normalized ARD count and Simpson’s diversity coefficient. The analysis yielded a mild positive correlation of 0.17, which, like the previous correlation, was deemed statistically insignificant due to its smaller critical *t* value. Although we did not find any statistically significant correlation between the two tested factors, namely the diversity of taxa and the age of samples, it is possible that spatio-temporal factors and the presence of particular groups of organisms could be responsible for the varied distribution of ARDs across different metagenomes.

### Selection pressure on ancient versus contemporary antibiotic inactivation enzymes

The evolutionary pressure, measured by the Ka/Ks ratio on proteins, is often assessed using the ratio of substitution rates between non-synonymous and synonymous mutations. Initially developed to evaluate selection pressure in distantly divergent proteins, this method is now being applied to phylogenetically closer proteins and even proteins from single populations. In the case of antibiotic resistance determinants (ARDs), enzymes capable of inactivating antibiotics, we aimed to determine the highest similarity percentage that would yield a divergence distance close to 1. Additionally, we sought to determine whether the popular Yang-Nielsen (YN) model or the more recent Model Averaged (MA) method in the Ka/Ks calculator would be optimal for identifying the best selection pressure ratio. To achieve this, we selected five pairs of genes ranging from 33% to 95% similarity and calculated their respective divergence distances and Ka/Ks ratios using both YN and MA methods. Notably, we included two 50% similarity sequence pairs, one from the same species and another from two different species. Our analysis revealed that the YN method could yield unrealistic divergence distances for ARDs and higher p-values compared to the MA method. Therefore, we concluded that the MA method was more suitable for our entire set of proteins. Additionally, we specifically chose proteins with similarity percentages equal to or less than 50% compared to the reference protein for Ka/Ks calculation.

All the tested antibiotic inactivation class ARDs were found to have Ka/Ks values well below 1, indicating that these genes experienced purifying selection. This suggests that the functions of these genes have been preserved, as expected for this class of genes. Notably, in project PRJEB47746, there was a diverse distribution of Ka/Ks ratios ranging from 0.01 to slightly above 0.5, as shown in Figure 5. Across all projects, antibiotic inactivation ARDs exhibited Ka/Ks values concentrated around 0.2, indicating strong negative selection pressure on these ARDs, as assessed by the MA method in the Ka/Ks calculator.

**Figure 5.**
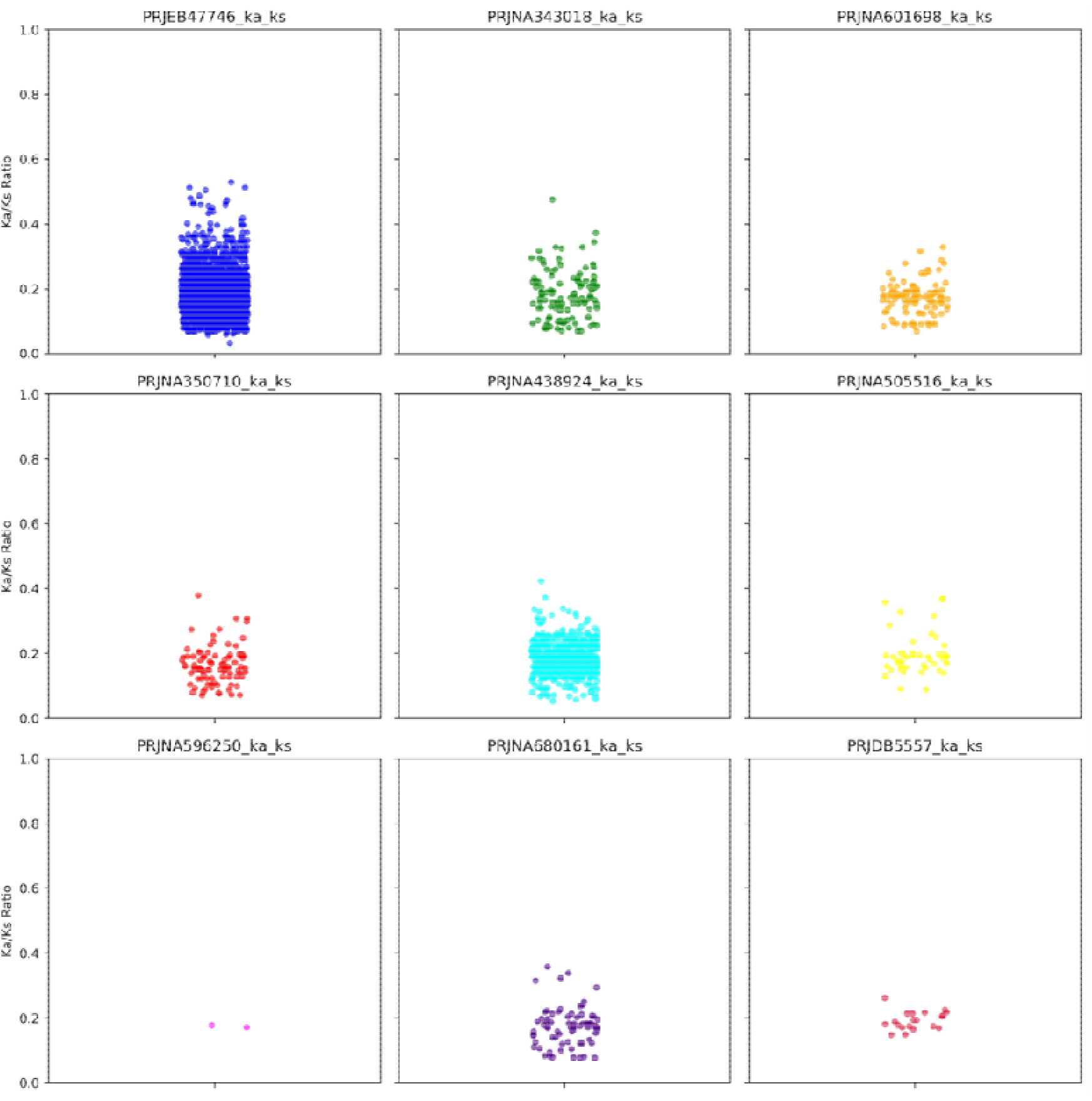
Distribution of Ka/Ks ratio for antibiotic inactivation genes from the ancient metagenome versus contemporary genes in the reference CARD database.

### Insights from mapping first assembly attempt

ARDs were identified from samples that did not yield any or had a low number of contigs, which were then subjected to mapping against reference ARG databases using the KMA aligner. No ARDs above the set threshold coverage of 70% were identified in these samples. In contrast, a total of 15 ribosomal proteins were identified above the set threshold coverage in all samples combined. Sample SRR13615822 did not yield any ARDs or ribosomal proteins due to low depth of coverage. Similarly, sample SRR8188256 did not yield any of these genes despite having substantial depth. Notably, this sample was derived from environmental DNA. Two other samples, SRR8188252 and SRR8188255, which likely originated from intracellular bacterial cells, had 8 and 1 ribosomal proteins, respectively, but no ARDs.

### Complete Viral genome retrieval and ARD profiling

In order to retrieve viral contigs from the metagenome and assess the presence of antimicrobial resistance determinants (ARDs) within them, a systematic approach was adopted. Initially, a hierarchical classification method was utilized to classify the contigs assembled from the metagenome. DeepVirFinder, employing a deep learning method, was first employed for a preliminary classification of the contigs, with a length cutoff set at 3000 bp. Contigs scoring a viral probability of at least 0.7 were further selected for classification. Subsequently, VIBRANT version 1.2.1 was employed for a more detailed classification, focusing on the identification of viral hallmark proteins within the contigs. Contigs classified as complete-circular by VIBRANT were then extracted and subjected to AMRFinderPlus analysis to profile ARDs within the complete viruses, employing a minimum identity and coverage threshold of 0.40 each. Through this comprehensive analysis, a total of 380 putative complete viruses were identified from seven projects (table 4) and none of these viruses were found to harbour ARDs.

**Table 4:** Complete Viral sequences identified by VIBRANT.

| Project Number | Number of complete viral genomes | Smallest genome size | Largest genome size | Average genome size |
| --- | --- | --- | --- | --- |
| PRJEB47746 | 282 | 3139 | 175833 | 31103 |
| PRJNA680161 | 13 | 3940 | 103189 | 49413 |
| PRJNA350710 | 16 | 3996 | 7581 | 5487.1 |
| PRJNA601698 | 19 | 3053 | 69374 | 20162.4 |
| PRJNA343018 | 6 | 5513 | 5513 | 5513 |
| PRJNA438924 | 38 | 3322 | 114447 | 43895.2 |
| PRJNA505516 | 6 | 5513 | 42098 | 11610.5 |

## Discussion

Our study encompassed a total of 2.3 tera base pairs of data, either assembled or directly profiled for ARD prevalence using prominent tools and databases such as AMRFinderPlus, KMA, and the CARD and RefGene catalogues of Antibiotic Resistance Genes. Approximately 1.3 billion contigs, each exceeding 1000 bps, were generated through the assembly process. However, four sets of raw metagenome reads from two distinct projects could not be assembled into contigs.

The AMRFinder analysis identified about 33,000 ARDs across all projects, meeting the criteria of at least 0.90 coverage and 0.40 identity with the reference database. To ensure accurate selection pressure analysis downstream, a coverage threshold of 0.90 was set, allowing only BLASTX results to remain while genes identified as putative ARDs by PARTIALX were removed. Additionally, ARDs with internal stop codons were included. Previous research highlighted the prevalence of divergent ARDs with low identity thresholds, potentially indicating phenotypic antimicrobial resistance characteristics[3]. To mitigate false positives, the presence of conserved domains in putative ARDs was verified using InterPro search.

The relative abundance of ARDs in ancient microbiomes was assessed by profiling ubiquitously present ribosomal genes. It was found that there were at least 2 ARDs for every identified ribosomal gene (5s, 16s, and 23s collectively) in the ancient metagenome. Notably, the relativistic estimate varied greatly among different projects including contemporary studies.

The distribution of identity percentages of ARDs from all samples was plotted into a box and whiskers plot, revealing a prevalence of ARDs around 40% similarity regardless of the microbiome’s temporal status. However, municipal waste and landfill metagenomes exhibited a higher likelihood of finding ARDs above 0.70 compared to pristine counterparts. It is noteworthy that the majority of ARDs were identified below 60% identity in both the ancient microbiomes and the contemporary microbiomes. But just mere detection of ARDs does not mean they would be expressed phenotypically, it can only be verified experimentaly.

The classification of ARDs into broad categories—antibiotic inactivation, antibiotic efflux, or antibiotic target alteration/protection/replacement—was enabled by the CARD Resistance Gene Identifier tool. Interestingly, while ancient microbiomes showed fewer inactivation-type ARDs, contemporary metagenomes displayed a more even distribution among the three classifications.

Further investigation aimed to understand the correlation between ARD prevalence and factors such as age and diversity. Initially, we investigated whether age influences the occurrence of ARDs. Utilizing an inverse-ranked Spearman correlation analysis, we found that age had an insignificant effect on ARD prevalence. Subsequently, we examined whether diversity correlates with ARD presence. Taxonomy was assigned to contigs using the Kraken2 tool, and Simpson diversity indices were estimated. Pearson correlation coefficients were then calculated between diversity indices and ARD abundance. This analysis also revealed an insignificant correlation between ARDs and the diversity of organisms present. In project PRJEB47746, a diverse range of contigs were found to carry ARDs, whereas in project PRJDB5557, only a few specific taxa were associated with ARDs.

Interestingly, thermophiles were present in all projects except PRJDB5557. These metagenomes, sourced from the Chuckochi region, likely experienced volcanic activity around 10 million years ago, potentially leading to the presence of active hot springs long after the landmass formation. Consequently, taxa such as *Aquificae, Thermomicrobiota*, and *Deinococcota*, which thrive in thermophilic environments, were identified. In contrast, PRJDB5557 samples were collected from the Tien Shan Mountain range, formed by the collision of Eurasian tectonic plates.

For our selection pressure estimation, we focused solely on antibiotic inactivation-type ARDs. This decision was made because even single nucleotide polymorphisms (SNPs) can affect the functionality of the resistance determinants in the other two categories, which do not require substantial evolutionary modifications. We calculated the ratio of non-synonymous mutations to synonymous mutations and found that all the antibiotic inactivation ARDs exhibited purifying selection. It is not surprising that the functionality of such enzymes is under negative selection pressure. Additionally, the timeframe tested here is relatively short in comparison to the evolutionary scale of microorganisms, if we analyse further ancient sequence there is a possibility of finding positive selection pressure. This finding is consistent with other studies reporting purifying selection in ARDs overall[31]. However, it’s important to note that many studies estimate selection pressure between closely related genes or those with considerably higher similarity, which can result in confounding estimates of selection pressure. Similar sequences typically exhibit lower substitution rates, making estimates derived from such low rates statistically unreliable [31,32]. Thus, we chose 50% identity as a cutoff below which the divergence would be sufficient to enable ka/ks analysis between them.

In our study, for samples that could not be assembled into contigs, we directly mapped the adapter-removed reads to the ARD database using the KMA aligner. However, even with this approach, we did not identify any ARDs in these samples. Additionally, it appears challenging to identify any genes from these reads, similar to the observation with small subunit ribosomal genes (see Table 3). Interestingly, two of these samples (labelled ‘i’ in superscript, Table 3) possibly harboured thriving organisms according to the original study.

Notably, from the contigs classified by VIBRANT from the metagenome-assembled contigs, no ARDs were found above the set threshold. Several reports claim that bacteriophages carry ARGs, while few report contrary findings. According to our literature review, we believe that bacteriophages typically do not carry ARGs, but they do so rarely[33]. Additionally, we learned that some specialized bacteriophages, known as phage-plasmids, can harbour ARGs. We sought to understand the scenario ages ago regarding bacteriophages retrieved from ancient permafrost. However, based on the limited results obtained in our study, it is too early to conclude whether ARDs are being carried in a few phages due to pressure from antibiotic usage in the modern times since there are no ARDs identified in the putative viral genomes from the ancient microbiome.

Based on these findings, it was concluded that the distribution of identity percentages between contemporary and ancient metagenome-sourced ARDs remained similar. Additionally, antibiotic inactivation-type ARDs from ancient microbiomes did not seem to have undergone substantial evolutionary changes to become more effective against antimicrobials in the antibiotic era. Importantly, none of the identified putative viruses carried ARDs in ancient times, although further investigation is warranted to see if the presence of few ARDs in the contemporary phage genomes are due to selection pressure from irrational use of antibiotics.

## Conclusion

The popular maxim “Everything is everywhere, but environment selects” appears to hold true to some extent concerning antibiotic resistance determinants. Antibiotic resistance has always existed and will continue to do so, as it is inherently encoded in the genomes of microbes. However, it is evident that anthropogenic influences have significantly exacerbated the effects of antibiotic resistance. Although these genes have been subject to negative evolutionary selection, they may exhibit greater phenotypic expression in contemporary times compared to the pre-antibiotic era.

## Data availability

All the metagenomic raw data were retrieved from NCBI SRA with the following accessions: PRJEB47746, PRJNA266334, PRJNA343018, PRJNA350710, PRJNA505516, PRJNA596250, PRJNA601698, PRJNA680161, PRJNA438924, PRJDB5557, PRJEB28336, PRJNA718480 and PRJNA606662. Any other additional data generated during the analysis shall be made available upon request. All the tools and software used are open-sourced.

## Acknowledgement

The authors express gratitude to the KBase platform funded by US Department of Energy, without which the assembly of some metagenome samples in this study would be impossible. Apart from that, S Gomathinayagam would like to thank ICMR for its fellowship support.

